# Transcription factor co-binding patterns drive conserved regulatory outcomes

**DOI:** 10.1101/189571

**Authors:** Adam G. Diehl, Alan P. Boyle

## Abstract

The mouse has been widely used as a model system in which to study human genetic mechanisms. However, part of the difficulty in translating findings from mouse is that, despite high levels of gene conservation, regulatory control networks between human and mouse have been extensively rewired. To understand common themes of regulatory control we look beyond physical sharing of regulatory sequence, where extensive turnover of individual transcription factor binding sites complicates cross-species prediction of specific functions, and instead look at conserved properties of the regulatory code itself. We define regulatory conservation in terms of a grammar with shared, species-specific, and tissue-specific segments, and show that this grammar is more predictive of shared chromatin states and gene expression profiles than shared occupancy alone. Furthermore, we demonstrate a marked enrichment of disease associated variation in conserved grammatical patterns. These findings offer new understanding of transcriptional regulatory mechanisms shared between human and mouse.

## BACKGROUND

As genome-wide datasets, including DNase hypersensitivity, transcription factor (TF) binding, and chromatin modifications, accumulate across multiple tissues and species, categorizing and ascribing functions to newly-identified putative regulatory sequences is a growing challenge. Traditionally, we have relied heavily on comparative genomics to make such predictions, either by extrapolating the function of a sequence based on known function of the orthologous sequence in another species, or by integrating data from multiple species to surmise the most likely function for a region. This strategy is effective within coding sequences, which have a fairly simple underlying grammar, retain relatively stable functions through evolutionary time, and are marked by extensive sequence conservation at the nucleotide level. However, while active regulatory elements are associated with a variety of characteristic functional markers (e.g., open chromatin and specific histone modifications), the underlying regulatory landscape is much more fluid between cells and species than for coding genes. Although we can use such measures to catalog putative regulatory regions across multiple species, detailed analyses of the underlying sequence conservation and effects on expression of nearby genes have shown limited correlation between positional conservation and regulatory output. Furthermore, we know that a large fraction of functional regulatory elements exist in sequences that are unique to individual species, such as transposable element (TE) insertions, which have been shown in multiple species to be frequently exapted for regulatory purposes. For example, it has been shown that TEs of the B2 SINE family have actively dispersed CTCF binding sites throughout the mouse genome [1], and that these B2 SINE insertions serve as anchor points for mouse-specific structural contacts [2]. It hardly needs to be stated that phylogenetic comparisons are not informative for these sequences.

An alternative approach to predicting the function of regulatory sequences is to focus not on the underlying conservation of a given regulatory locus, but rather the regulatory grammar expressed in the combinations of co-bound transcription factors, or cis-regulatory modules (CRMs) we observe in different species and tissues. Indeed, there is considerable interest in CRMs as drivers of adaptive evolution [3-6]. We know that most gene expression is mediated by combinations of TFs acting cooperatively to regulate their target genes [7], that distinct sets of TFs elicit specific and reproducible gene expression patterns [8-11], and that these regulatory outcomes often directly translate between species [12]. This suggests that a conserved regulatory grammar may underlie control of common physiological pathways, and that species-specific modifications to this grammar and/or how it is used may explain much of the complexity in gene expression programs among species. However, the evolution of this regulatory grammar has never been extensively analyzed, in part because of the inherent complexity of the regulatory landscape, given that there are ~1200 TFs predicted in mammals [13].

Because it is intractable to directly infer the functions of all possible combinations of TFs, we need to simplify the regulatory landscape. Much like in natural language, we expect that many of the regulatory sentences we observe in mammalian genomes will share equivalent meanings. As such, we can apply machine learning techniques to reduce the regulatory landscape into sets of significantly co-occurring combinations of TFs. Self-organizing maps (SOMs) are particularly well-suited for this purpose because of their ability to project such high-dimensional data onto a two-dimensional grid topology, facilitating easy comparisons of various properties of the data. SOMs have been previously used to describe CRM data in humans [7] and across distantly-related species [14]. Here we apply this method to sets of matched immune cells from human and mouse; as far as we are aware, this is the first application of an SOM to infer shared and species-specific regulatory properties across two mammalian species.

The mouse has long represented a flexible genetic system for exploring human genetic disease. However, because of the estimated 75 million years of divergence separating these two species [15], only ~40% of their DNA sequences are directly alignable. We know that protein coding genes remain largely unchanged while noncoding sequences, which harbor most of the gene regulatory circuitry, have undergone extensive turnover. We know that, even though the pool of TFs and their binding specificities have changed little among vertebrates, the majority of occupied TF binding sites (TFBSs) are species-specific, even in the same tissue among closely related vertebrates [16]. At the evolutionary distance between human and mouse, sequence turnover frequently changes the order, orientation, and position of TFBSs relative to their target genes even when regulatory outcomes show evidence of conservation [17]. Such sequence-level changes are likely to obscure conserved patterns of combinatorial TF binding, complicating the use of sequence conservation to elucidate the regulatory code. If we are better able to map the similarities and differences in regulatory control between species, we may be able to predict which genetic pathways retain similar properties and thus which findings from mouse are likely to translate back to the human system.

Recent advances in sequencing technologies have allowed us to begin mapping regulatory regions in human and mouse and started revealing the extent of turnover at regulatory sites in human and mouse. Early work has demonstrated that only 40-50% of human and mouse regulatory elements reside in orthologous sequence, and, at a majority of these sites, TF binding is not maintained between species [18,19]. However, the functional consequences of such changes are unclear. To shed the inherent constraints of methods based on shared-occupancy, we explore regulatory evolution by analyzing conserved combinatorial TF binding patterns independent of their sequence context. We do so by simultaneously comparing TF ChIP-seq co-binding profiles from four human and mouse cell types using a SOM, ignoring factors such as the order, orientation, spacing, cell, and species context. By doing so, we allow conserved and divergent properties of the regulatory code to emerge in the partitioning of co-binding profiles, or grammatical patterns, identified by the SOM. We demonstrate that much of the regulatory logic between these human and mouse cell types has remained intact despite extensive turnover at the level of individual TFBSs. By mapping additional annotations to the SOM map, we further show that grammatical patterns show stable functional signatures, are correlated with tissue-specific expression profiles of nearby genes, and are enriched for functional GWAS variants for relevant human phenotypes, making them potentially useful predictors of functional specificity and conservation.

## RESULTS

### Grammatical patterns are significantly more conserved than individual regulatory loci

We used a self-organizing map (SOM) to learn significantly occurring co-binding patterns, which we call grammatical patterns, for 27 general TFs among two sets of biologically matched cell types from human and mouse: K562 (K) and MEL (M) myeloid cells, and GM12878 (G) and CH12 (C) lymphoid cells (**Fig. 1A-C** and see Methods) [7]. ChIP-seq peak data for each of the factors in all four cell types were obtained from the ENCODE portal [19].

**Figure 1:**
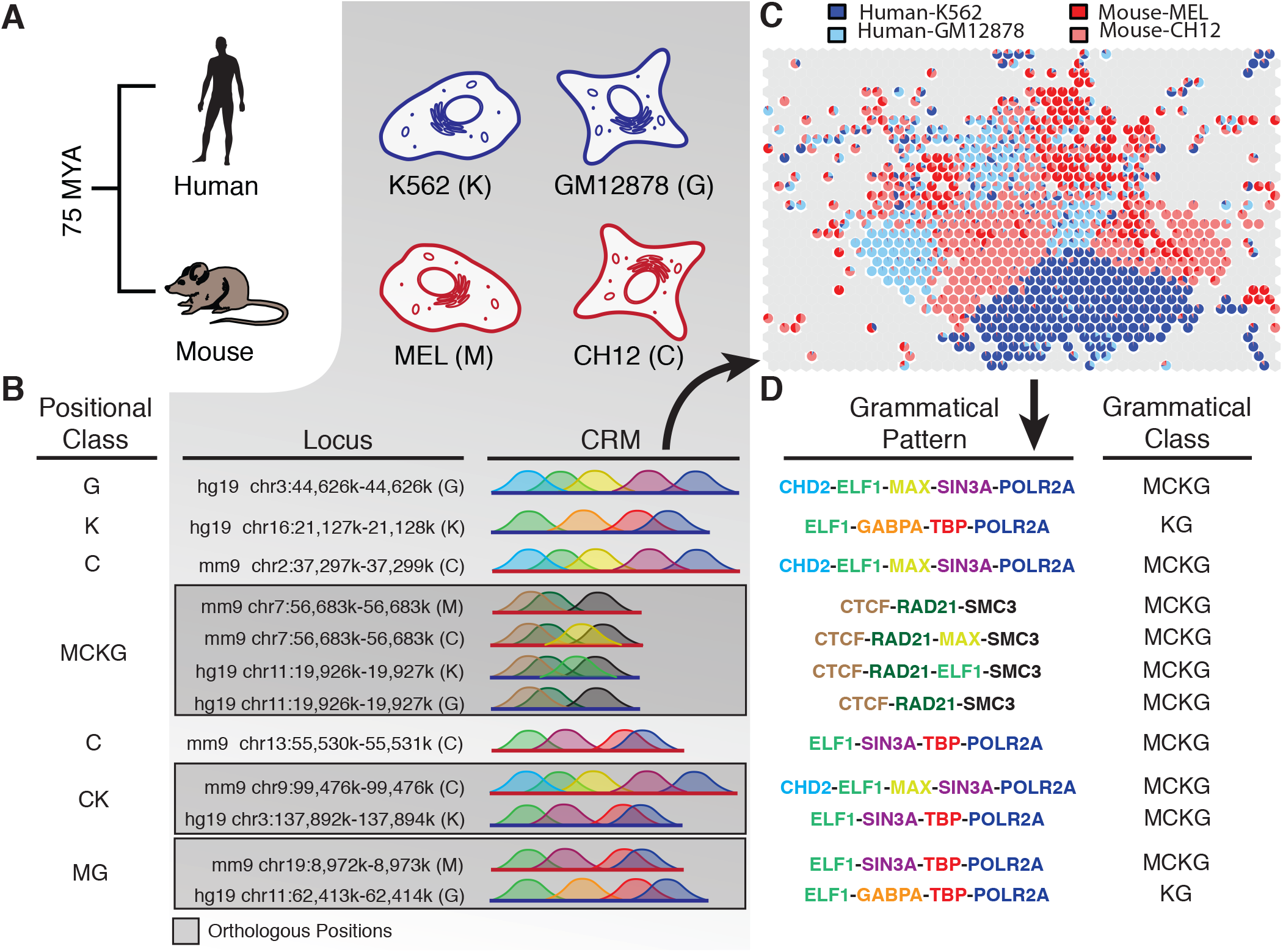
The self-organizing map, grammatical classes and positional classes. **(A)** ChIP-seq datasets from human and mouse myeloid (K562 and MEL) and lymphoid (GM12878 and CH12) immune cells were obtained from the ENCODE DCC portal. **(B)** Overlapping ChIP-seq peaks were assembled into cis-regulatory modules. The “Positional Class” table contains locus-level assignments of the example CRMs to their respective <u>positional classes</u>, which describe the set of cells in which we observed shared occupancy at a given locus. Previous studies have used shared occupancy and/or shared presence of open chromatin as a measure of regulatory conservation, and here we extend the concept of “positional conservation” introduced by Vierstra et al. [18], to include shared occupancy of a CRM at the same locus between species and/or tissues. **(C)** A self-organizing map (SOM) algorithm was used to infer co-occupancy patterns significantly overrepresented in these four cell types. After removing patterns observed in any of 10,000 permutations of the dataset, 780 co-occupancy patterns, which we call <u>grammatical patterns,</u> remained. These are shown as colored hexagons on the map**. (D)** Grammatical patterns are defined as unique co-occupancy profiles of two or more TFs. The “Grammatical Pattern” table shows the co-occupancy profile corresponding to the grammatical pattern to which each example CRM in (B) is assigned. Note that, although all example CRMs in (B) within the same grammatical class are shown as containing identical ChIP-seq profiles, this is not a requirement, and individual TFBSs within a CRMs may actually occur in any order, spacing and orientation within a CRM, and some flexibility is allowed in specific TF content. The “Grammatical Class” table shows the assigned grammatical class for each example CRM, which is defined by the set of cells in which its assigned grammatical pattern is observed at a rate significantly exceeding random expectation.

These were aggregated into cis-regulatory modules (CRMs) based on mutual overlap of peaks and encoded as binary occupancy vectors, which were assigned to grammatical patterns by the SOM algorithm. After filtering out all patterns that occurred in any of 10,000 random permutations of the data, 780 distinct grammatical patterns remained (**Fig. 1C** and **1D**). Grammatical patterns in our dataset reflected many known co-associations between TFs [7,14,20], and generally reflected previously-reported spatial biases for their member TFs [20], with most patterns biased toward either promoters or enhancers. Overall, 76% of shared grammatical patterns showed stable spatial preferences between human and mouse and, of the remaining 164 patterns, 159 contained ETS1, which is known to have switched in spatial preference from promoter-proximal regions in human to distal regions in mouse [20], or other TFs known to co-associate with ETS1 [20].

Each grammatical pattern was assigned to a grammatical class, based on the cell(s) in which a pattern was observed at a rate exceeding a random expectation (**Fig. 2A**). Grammatical classes reflect a pattern’s species/tissue-specificity and, by extension, its depth of conservation. That is, grammatical patterns that are used across multiple species and cell types likely serve general, and perhaps more deeply conserved, regulatory functions while those observed in a narrow range of cell types may serve tissue/species-specific or emergent functions.

**Figure 2:**
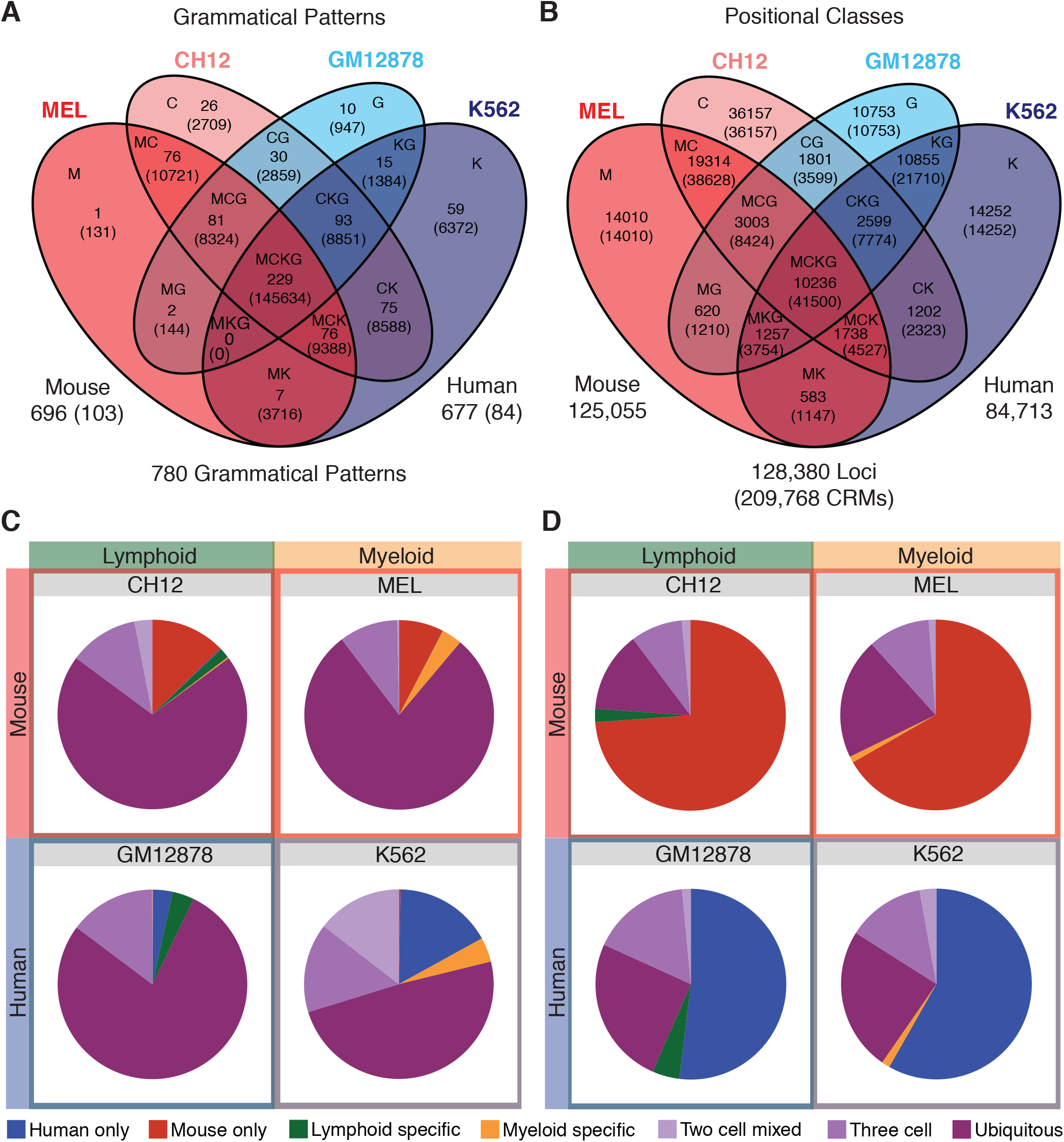
Grammatical patterns capture regulatory conservation better than shared occupancy. **(A-B)** Venn diagrams represent segmentation of regulatory space into 15 possible grammatical classes (A), and positional classes (B), defined by the cell(s)/species in which a grammatical pattern is used, or a regulatory locus is occupied. Class labels were assigned based on the first letter of the cell type(s) represented: (M)EL, (C)H12, (K)562, (G)M12878, to produce the set of labels: M, C, K, G, MC, MK, MG, CK, CG, KG, MCK, MCG, MKG, CKG, and MCKG. **(A)** Grammatical classes describe regulatory conservation as defined by distinct combinations of co-bound TFs shared across cells and/or species, and are assigned to grammatical patterns based on the set of cells in which they were observed. Each segment in the Venn diagram carries the grammatical class label, the number of grammatical patterns within the class (first number), and the number of CRMs among all patterns within the class (number in parentheses). Segments marked with a single cell label represent tissue/species-specific regulatory functions, which may be less conserved, while those spanning species and/or tissues are considered more conserved. Overall, the SOM partitions the dataset into 780 grammatical patterns, 593 of which are used in both species and 187 that are species-specific (103 mouse and 84 human). This represents a significant reduction in complexity compared to positional conservation. **(B)** Positional classes describe regulatory conservation in terms of shared sequence occupancy. We define a regulatory locus as a distinct location within one species and its orthologous locus (if present) in the other species. Regulatory loci were assigned to positional classes based on the cell(s) in which CRM occupancy was observed. Each segment is labeled with its corresponding positional class, the total number of loci within the class (first number), and the total number of CRMs assigned to the class (number in parentheses). In all, 209768 CRMs were observed at 128380 distinct loci, 54491 of which are occupied in both species. Whereas grammatical conservation concentrates most loci into shared and ubiquitous grammatical classes, positional conservation spreads loci more broadly across shared, species-specific, and cell-specific classes. Notably, 58% of all loci in (B) are occupied in only one of four cell types, whereas only 5% of CRMs in (A) carry grammatical patterns classified as cell-specific. **(C-D)** Grammatical patterns capture more regulatory conservation than positional conservation as shown by a substantially higher fraction of CRMs assigned to shared aggregate classes. Pie charts denote the fraction of CRMs in each cell type assigned to aggregate classes defined as human-only (K, G, KG), mouse-only (M, C, MC), lymphoid-specific (CG), myeloid-specific (MK), two-cell-mixed (MG, CK), three-cell (MCG, MCK, MKG, CKG) or ubiquitous (MCKG) based on their assigned grammatical **(C)** or positional **(D)** class.

Recent studies have shown positive correlations between shared TF occupancy and functional outcomes including tissue-specific enhancer activity [20] and tissue-specific expression of nearby genes [20,21]. However, while these studies and others [22,23] suggest a positive correlation between deeply-shared orthologous TF binding and conserved regulatory outcomes, they focus on tissue-specific TFs and thus their findings may not generalize well to broader tissue sets. By contrast, there is substantial evidence for extensive tissue-specific repurposing of positionally-conserved regulatory elements, with up to 69% of DHS shared between human and mouse repurposed across species [18]. It seems likely that this repurposing is based, at least in part, on changes in TF binding across species, given that multiple studies have shown that only a minority of CRMs and DHS shared between human and mouse also share equivalent sets of TFs [17,18,22-25]. In support of this hypothesis, Vierstra et al. also report that positional and operational TF binding conservation were strongly enriched in DHS with shared tissue activity while tissue-specific and cell fate-modifying TFs are specifically depleted in DHS that had been repurposed into a new tissue. These findings strongly suggest that the TF co-binding profile occupying a regulatory locus directly stipulates its function, independently of whether it is also active in other tissues/species. Notably, Vierstra et al. also found a corresponding enrichment of tissue-specific TFs in DHS that retained tissue-specific activity across species, strengthening the link between conserved regulatory grammar and conservation of function. Accordingly, it seems likely that changes in regulatory grammar are an important agent in regulatory evolution, and that regulatory grammar can be useful in predicting conserved and divergent regulatory sequence function.

Here we extend the concept of positional conservation, first introduced by Vierstra et al. [18], to describe shared occupancy of a CRM in the orthologous locus across species and/or tissues, regardless of the specific TFs co-bound therein. To evaluate the extent to which positional conservation is correlated with regulatory grammar in our dataset, we cross-mapped CRMs to orthologous regions of the human/mouse genome with bnMapper [24], grouped them into distinct orthologous and non-orthologous loci, and assigned each locus to a positional class based on the cell(s) in which a CRM was present at the given genomic location (**Fig. 2B**).

We began by characterizing the degree of tissue-specific repurposing of shared CRMs between tissues and species. Consistent with previous studies [17,18,21,23], we found that only 30-50% of CRMs were alignable between human and mouse (**Fig. 2D**). At these shared loci, we interpreted a change in grammatical pattern between cells as evidence for repurposing, which we observed at 74% of positionally-conserved loci (**fig. S1**). Interestingly, the fraction of repurposed loci was positively correlated with the depth of shared occupancy, and >90% of loci in the MCKG (occupied in all four cell types) positional class showed evidence of repurposing in at least one cell. These findings corroborate previous observations of extensive TFBS turnover between orthologous regulatory loci at all evolutionary distances [17,22,23,26,27], suggesting that positional conservation likely has limited predictive value for determining the output of a regulatory sequence.

Despite the high rates of TFBS turnover we and others observe at orthologous regulatory loci, previous studies have suggested that tissue-specific regulatory logic, at the level of TF co-association, is at least partially conserved across species, even at large evolutionary distances [14,20,21,25]. Indeed, we show that 89% of CRMs host grammatical patterns shared between human and mouse (**Fig. 2C**). Furthermore, we saw no clear association between grammatical class membership and evolutionary history of a CRM, with shared grammatical patterns contributing 88% of orthologous CRMs and 92% of non-orthologous CRMs. Therefore, although individual regulatory loci may be frequently repurposed, most of the TF co-binding patterns found in human and mouse appear to be deeply conserved. This suggests that the underlying regulatory logic by which specific regulatory outcomes are produced is also deeply conserved. Taken in context with previously reported findings, these results suggest that grammatical patterns likely reflect specific regulatory functions, and by extension, functional conservation, more closely than phylogenetic conservation. As such, grammatical patterns may be a useful tool for inferring whether positionally-conserved CRMs retain equivalent functions across species, as well as inferring the regulatory activities of non-orthologous CRMs, for which phylogenetic clues to their potential functions do not exist.

### Grammatical classes have stable functional signatures across orthologous and species-specific CRMs despite differential sequence conservation

Given that only a minority of human and mouse CRMs are alignable between human and mouse, the remaining 50-70% of CRMs cannot be functionally classified using phylogenetic comparisons. In order to further evaluate the ability of grammatical patterns to functionally characterize these sequences, we divided non-orthologous CRMs into species-specific gain and loss fractions by cross-mapping to three outgroup species: dog, horse, and elephant, and applying a phylogenetic maximum parsimony algorithm to determine the most likely branch on which a sequence was gained or lost. To examine the degree of correlation between functional chromatin signatures and phylogenetic conservation, we measured intersections between DHS, active chromatin states (ACS) from ChromHMM [28], and PhastCons elements (PE) [29], within orthologous, gain, and loss fractions of the dataset across six aggregate grammatical classes: human-specific (K, G, KG), mouse-specific (M, C, MC), lineage-specific (MK, CG), two-cell-mixed (MG, CK), three-cell (MCG, MCK, MKG, CKG), and MCKG.

DHS, which are commonly used to indicate noncoding regulatory function [30], had remarkably consistent intersections between orthologs, gains, and losses (**Fig. 3A, table S3**). All aggregate classes of CRMs were enriched for DHS relative to matched background sequences (all p-values < 2.1e-24, table S3) and, in total, 88% of CRMs overlapped DHS. Similarly, the percentage of CRMs labeled with ACS remained relatively stable between orthologous, gain, and loss fractions in all aggregate classes (Fig. 3B), although the effect was not as uniform as with DHS, and all subsets but one were enriched for ACS in CRMs compared to matched background sequences (all p-values < 2.6e-3, table S4). Similar to previous observations [20], ACS in our dataset were present in 58% of all CRMs. For both DHS and ACS, we see a decreased intersection in the MCKG class in all three fractions. This effect can be attributed largely to cohesin-related grammatical patterns, and removing the two most prevalent cohesin-related patterns markedly reduces its magnitude (fig. S4), increasing intersection with DHS to 96% and ACS to 79%.

**Figure 3:**
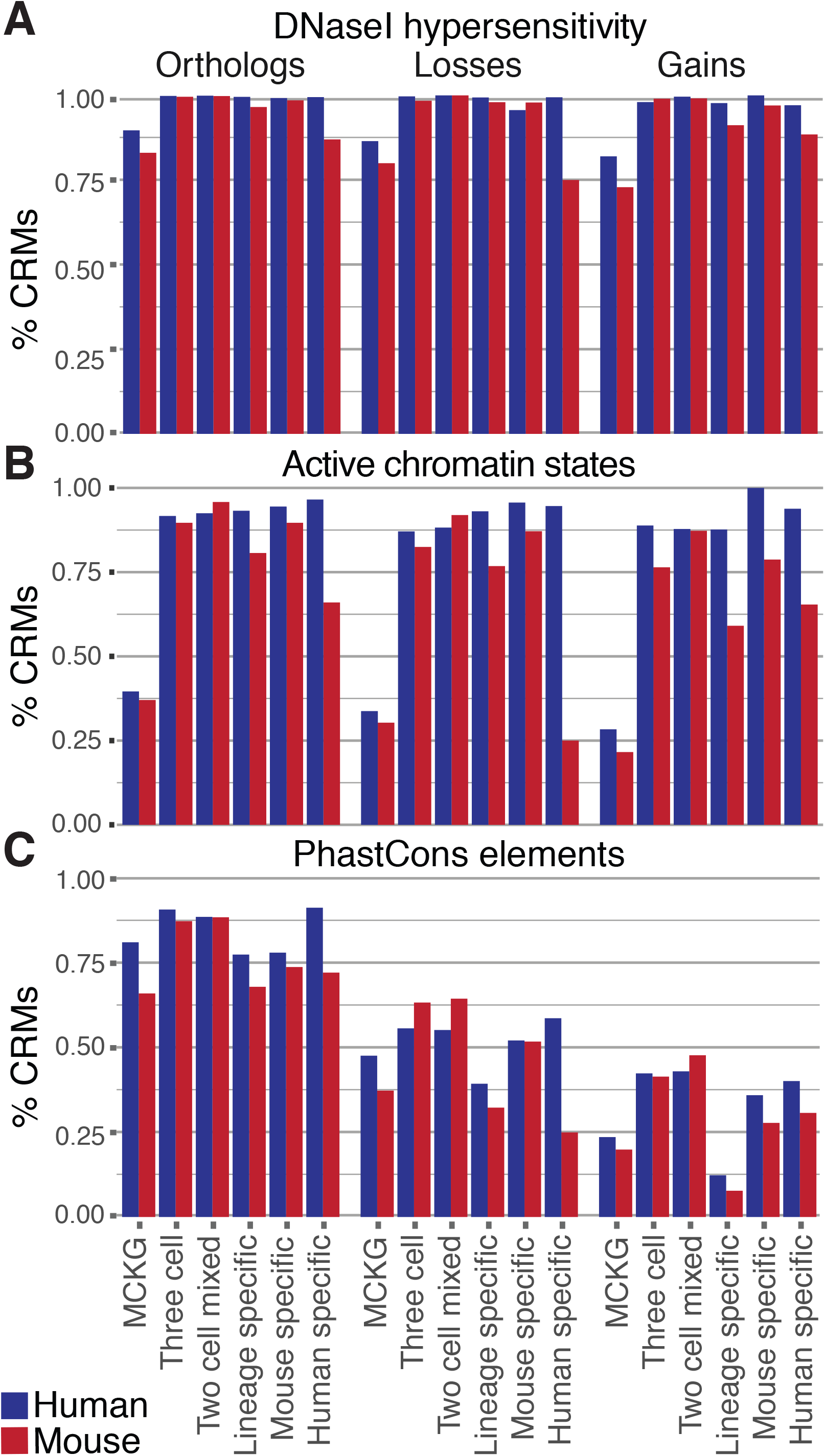
Grammatical patterns associate stably with genomic annotations associated with regulatory function, regardless of underlying sequence conservation. CRMs within six aggregate grammatical classes were divided into orthologous, and species-specific loss and gain fractions, based on presence/absence of orthologous sequence in human and mouse, and in three outgroup species. Within each subset, we quantified the fraction of CRMs overlapping DNaseI hypersensitive sites **(A)**, ChromHMM active chromatin states **(B)**, and PhastCons elements **(C)**. Whereas the overlaps with DNaseI hypersensitive site and active chromatin states, both of which are frequently used as a proxy for regulatory function, remain relatively stable among all subsets of the data (A-B), overlaps with PhastCons elements, which specifically measure phylogenetic conservation, decrease predictably between orthologous, loss, and gain CRMs (C).

In contrast to the stability in functional signatures we observed with DHS and ACS, intersection between CRMs and PE, which reflect functional constraint measurable by phylogenetic conservation, declined steadily between orthologous, loss, and gain fractions of the dataset (Fig. 3C). Overall, 65% of CRMs (58% mouse, 75% human) contain PE and CRMs were 3.3 times more likely to contain PE than matched background sequences (p < 6.2e-258). Excluding the two cohesin grammatical patterns only yielded a modest increase in PE intersection, to 68% of all CRMs. Significant enrichments over background were observed in all six aggregate classes among orthologs, gains and losses, except in three subsets with insufficient data (table S2). As expected, orthologous sequences contained systematically more PE than both losses and gains, and sequence conservation was correlated with the evolutionary age of the CRMs. The youngest elements, species-specific gains, showed the least overlap with PE, while PE intersection in losses occurred at an intermediate level relative to gains and orthologs (p-values < 2.2e-240). Notably, evolutionary conservation in many CRMs appears to arise largely from direct TF binding, as shown by clustering of phastCons scores approaching 1 within 50bp of ChIP-seq peak summits, and we see the same correlation between evolutionary age and phastCons scores near ChIP-seq peak summits evident in Fig. 3C (fig. S4). Based on the prevalence of active functional annotations shown by us within these regions, it follows that PE and, by extension, phylogenetic comparisons in general, are a poor indicator of regulatory function in these regions. By contrast, the close correlation we observe between grammatical conservation, DHS, and ACS, regardless of evolutionary history, is consistent with the hypothesis that grammatical patterns are generally predictive of regulatory potential.

### Grammatical patterns predict underlying chromatin states and grammatical classes are correlated with cell-specific chromatin state conservation patterns

To expand on the association between grammatical patterns and underlying ACS, we compared the ability of matched ACS to predict the grammatical patterns at pairs of positionally-conserved loci, and of matched grammatical patterns between pairs of loci to predict underlying ACS. We found that pairs of loci with matched grammatical patterns also had matched ACS 29% of the time (Fig. 4A) - a ~19-fold increase in predictive power over positionally-conserved loci with matched ACS, among which only 1.5% of pairs of shared the same grammatical pattern across cell types (Fig. 4B). This observation was stable between promoters, strong enhancers, and weak enhancers, although predictive power was lower in strong enhancers for both matched ACS and matched grammatical patterns (fig. S5A-B). The predictive value of grammatical patterns regarding underlying ACS can readily be seen in fine scale by plotting the chromatin states present for the CRMs within a grammatical pattern (fig. S5C). These findings are consistent with our expectation that grammatical patterns represent collections of CRMs with equivalent regulatory functions and, as such, are more predictive of cognate functional chromatin annotations than such marks are of latent regulatory logic.

**Figure 4:**
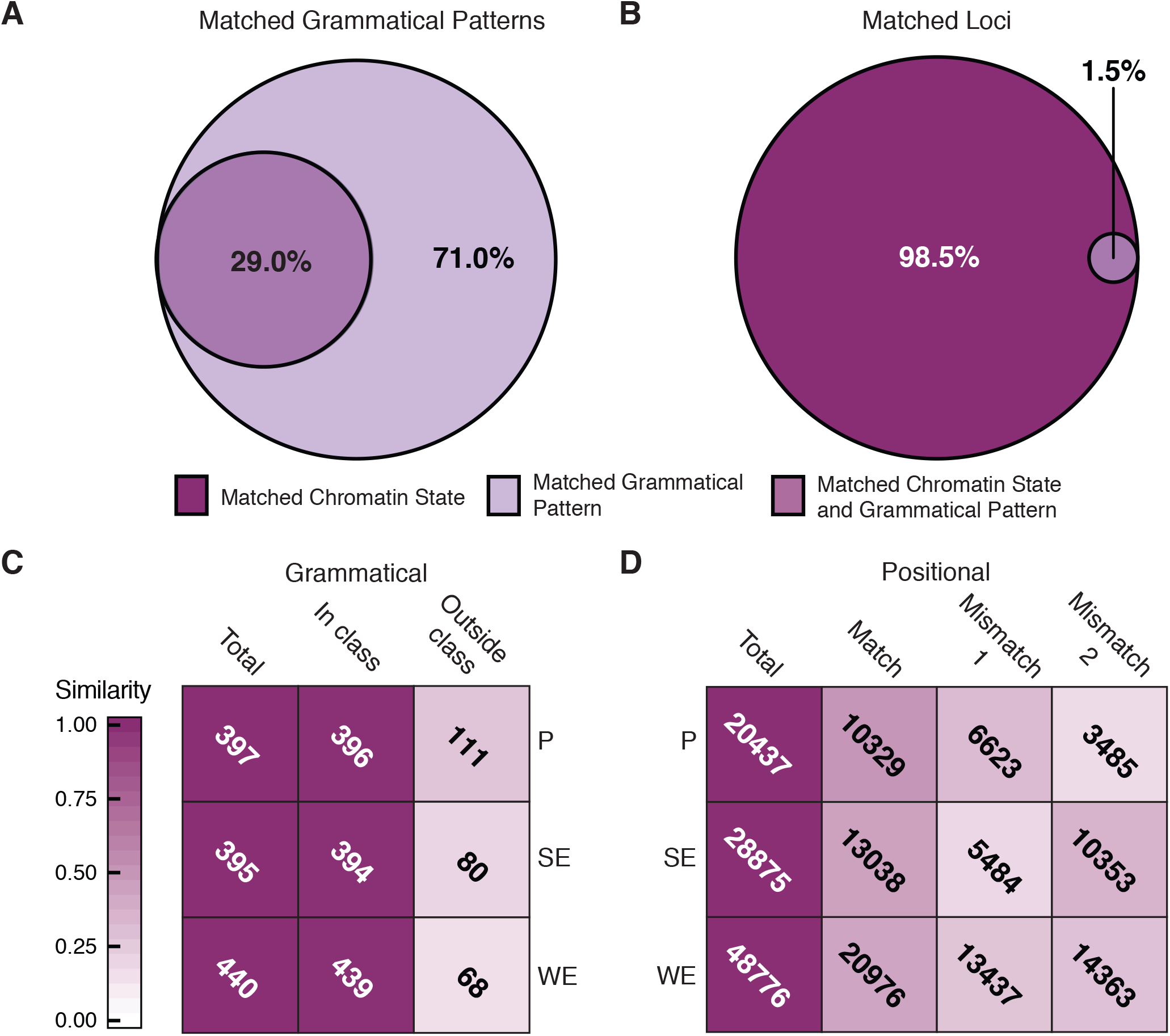
Grammatical patterns are more predictive of active chromatin states than positional conservation. We intersected all CRMs with predicted promoter (P), strong enhancer (SE), and weak enhancer (WE) chromatin states from ChromHMM. **(A)** To evaluate the predictive value of grammatical patterns for underlying chromatin states, we calculated the fraction of CRM pairs within each grammatical pattern with matching chromatin state annotations, which comprised 29% of all pairs. **(B)** By contrast, pairs of positionally-conserved sequences at the same/orthologous locus with matching chromatin states only carried matching grammatical patterns in 1.5% of cases, showing that positional conservation of chromatin states has limited predictive value for specific regulatory content. **(C-D)** Heat maps describe the extent to which grammatical classes (C) and positional classes (D) predict cell/species-specificity of underlying chromatin states. Shading density reflects the fraction of loci or patterns that carry the given chromatin state marks in the expected cell type(s) based on their assigned class. **(C)** All grammatical patterns in which at least 50% of CRMs intersect a given chromatin state were counted (“total” column). The “in class” column describes the number of patterns in each row in which at least 50% of CRMs from cell types belonging to its grammatical class carry the same mark (see fig S3B). The “outside class” column shows the number of patterns in each row in which the given chromatin mark is present in cell types not belonging to its grammatical class. **(D)** Chromatin states in all four cell types were determined for each locus in the dataset to evaluate the extent of agreement between positional classes and cell-specificity of chromatin marks at positionally-conserved loci. Loci carrying a given state annotation were counted (“total” column), and the number of loci from each row where the given state was observed in all cells belonging to its positional class, and no cells outside its class, were counted as matches (match column). Loci at which the given state was observed in cells outside the given class are counted in the “mismatch 1” column, while those lacking the state in at least one cell within the given class are counted in the “mismatch 2) column.

Given that grammatical patterns belong to distinct, cell/species-specific grammatical classes, we wanted to test our expectation that these also correspond to cell/species-specific functional classes. If our hypothesis is correct, we would surmise that ACS would match between all/most CRMs from cells belonging to a given grammatical class, while CRMs from cells that do not belong to a grammatical class (despite having been assigned to the given grammatical pattern) would be less likely to share the “correct” ACS. We tested this hypothesis by counting the proportion of grammatical patterns in which at least 50% of member CRMs from cell types belonging to its grammatical class contained a given ACS while a majority of member CRMs from other cell types lacked a matching ACS (fig. S6A). In all, 694 of the 780 grammatical patterns showed evidence of ACS specificity, according to the 50% threshold. In accordance with our hypothesis, ACS annotations within member cells closely corresponded to the grammatical classes of most patterns, with a remarkable 99.8% match rate among cells belonging to the parent grammatical class (Fig. 4C, fig. S7A), while only 15-28% of patterns had matching ACS in CRMs from cells outside the grammatical class. This level of specificity remained unchanged even with increasing thresholds for chromatin state specificity up to 100%, although the total number of patterns meeting the inclusion criterion decreased steadily as the threshold increased (fig. S8A-C). Interestingly, we see this class-specific pattern of ACS enrichment only when comparisons are referenced to cells belonging to a given grammatical class, but not when nonmember cells are used as the reference (fig. S7A). This phenomenon was not observed among positional classes (fig. S7B). Beyond simply demonstrating the predictive value of grammatical patterns regarding underlying ACS, these data also suggest that most grammatical patterns are active in different chromatin components, which may be expected based on the spatial biases we observed for most grammatical patterns.

For comparison, we asked how often positional classes matched underlying ACS conservation: i.e., that all occupied sequences share a given ACS while unoccupied sequences lack a matching ACS annotation. To determine if this is true, we counted the proportion of positionally-conserved loci at which all cells belonging to its positional class share a given ACS, while no cells outside the positional class carry the matching ACS (fig. S6B). To be considered a match, both of these criteria had to be met; loci failing to meet one or both criteria were classified into two mismatch categories: “mismatch 1”, where the same ACS was present in nonmember cell(s), and “mismatch 2”, where one or more member cells lacked the ACS. Notably, the “match” rates we observed at positionally-conserved loci were much lower that for grammatical patterns: 50% at promoters, 45% at strong enhancers and 43% at weak enhancers which is consistent with previous studies (Fig. 4D, fig. S7B) [19]. Relaxing the requirement for strict matching between all occupied cells at a locus yielded only modest increased in match rate (fig. S8D-E). This suggests that shared occupancy, in addition to being a poor predictor of grammatical patterns, also has limited predictive value in terms of conservation of functional chromatin annotations when compared to specific regulatory content. This can be easily seen in a plot of 10 randomly selected loci from the CG class, showing discordance between the positional classes, implied by occupancy, and ACS in 8 of 10 cases (fig. S5D). By contrast, grammatical patterns appear to be informative in terms of both categorical regulatory activities, as captured by their correlations with specific ACS, and of the depth of functional conservation of their underlying regulatory grammar, as expressed by the cell-specificity of their cognate ACS.

### Tissue and species-specificity of CRMs and grammatical classes correlate significantly with gene expression patterns

To expand on our finding that grammatical classes are associated with cell-specificity of underlying ACS, we wanted to see if there was also an association between grammatical classes and cell-specific gene expression patterns. If this is true, we would expect presence of tissue/species-specific CRMs to increase the likelihood of a matching tissue/species-specific expression profile in nearby genes. To test this theory, we looked for enrichments of tissue/species-specific grammatical patterns among CRMs nearest to matching tissue/species-specific genes (in-class genes), and depletion of the same patterns among CRMs nearest to genes with non-matching tissue/species-specific expression (cross-class genes), relative to stably-expressed genes.

We first generated lists of stably-expressed genes, and genes specific to human, mouse, lymphoid cells, and myeloid cells, using DESeq [31]. Within each of the corresponding aggregate grammatical classes, we calculated the fractions of CRMs nearest to genes with stable expression (πS) and CRMs nearest to in-class genes (πD) and calculated a tissue-specificity coefficient, *c*, as *c* = log2(πD/πS), where c > 0 indicates tissue-specific enrichment and *c* < 0 indicates tissue-specific depletion. We also calculated a corresponding coefficient, c′, as c′ = log2(πD′ / πS), where (πD′) represents the fraction of CRMs nearest to cross-class genes.

Consistent with our hypothesis, we observed significant enrichment of tissue/species-specific CRMs near in-class genes among all aggregate grammatical classes (Fig. 5, light purple bars). Likewise, we saw a consistent trend toward depletion of CRMs near cross-class genes, which was significant in all but one aggregate grammatical class (Fig. 5, dark purple bars). We also observed a modest but significant depletion of tissue/species-specific CRMs near stably-expressed genes (p = 3.4e-4). Notably, MCKG patterns were neither enriched nor depleted among CRMs near any category of genes. This is consistent with our theory that MCKG grammatical patterns serve broad regulatory roles spanning both tissues and species, while tissue/species-specific grammatical patterns are important in producing corresponding tissue/species-specific gene expression profiles.

**Figure 5:**
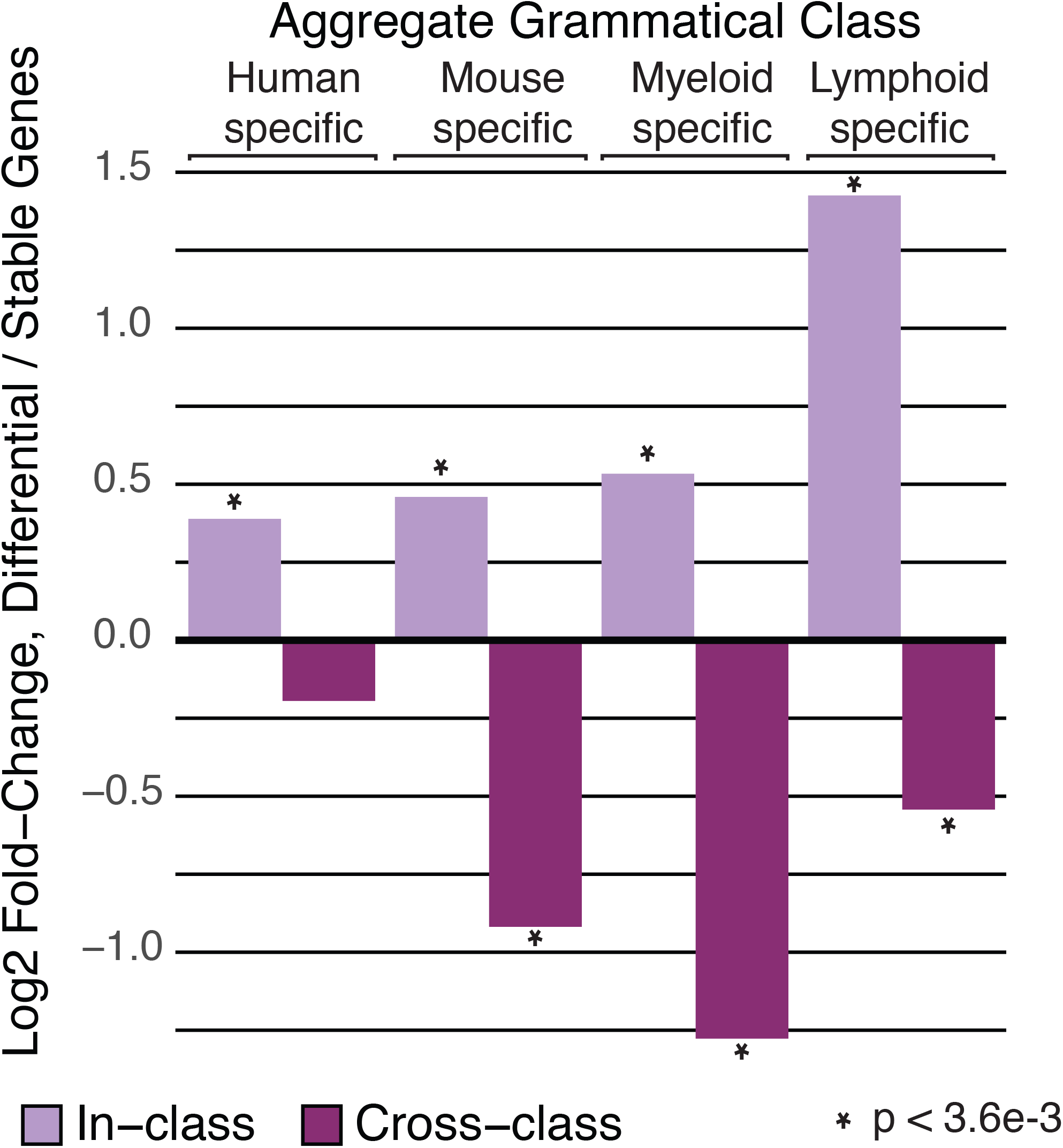
Gene expression patterns are correlated with cell- and species-specific grammatical patterns. To evaluate a potential causal relationship between CRMs within tissue-and species-specific grammatical classes and corresponding gene expression patterns, we calculated log-ratios of the proportion of tissue/species-specific CRMs targeting differentially-expressed genes compared to stably-expressed genes for each aggregate grammatical class. Light purple bars represent the log-ratio observed for tissue/species-specific CRMs targeting genes with the matching tissue/species-specific expression profile. Dark purple bars indicate log-ratios observed for tissue/species-specific CRMs targeting genes with mismatched tissue/species-specific expression profile. Positive values indicate enrichments while negative values indicate depletions. Statistical significance of enrichments/depletions was evaluated using Fisher’s exact test, with significant p-values indicated by a star. Notably, significant enrichment among matching tissue/species-specific genes was seen for all aggregate grammatical classes, along with corresponding depletions among nonmatching genes.

Based on the previous observation that mouse regulatory sequences without human orthologs are enriched near immune-specific genes [19], we wondered if we would see a similar enrichments of mouse-specific, non-orthologous CRMs near genes with mouse-specific expression, and of human-specific CRMs near human-specific genes. Using the same lists of species-specific and stably-expressed genes, we counted species-specific CRMs and CRMs with 1:1 orthologs nearest to species-specific genes and performed Fisher’s exact tests for enrichment versus stably-expressed genes. As expected, we found strong enrichments of species-specific CRMs near genes with expression patterns specific to the same species (Mouse: OR 2.1, p 3.3e-38; Human: OR 2.9, p 1.2e-69).

We observe that species-specific gene expression is correlated with enrichments of both species-specific grammatical patterns, and species-specific non-orthologous CRMs. This suggests that expression divergence is driven by a combination of TFBS turnover leading to creation of species-specific grammatical patterns at shared regulatory loci, and physical sequence gain/loss, creating regulatory loci unique to a species, and which we have shown predominantly carry conserved grammatical patterns. From these data, it is difficult to tell which effect is more influential, and it is likely that a complex interplay between both processes underlies species-specific divergence in target gene expression.

### Conserved and tissue-specific grammatical patterns are significantly enriched for tissue-appropriate human GWAS variants

If the grammatical patterns in our dataset are indeed important for proper gene regulation in immune cells, we would expect to find enrichments for immune-associated GWAS SNPs among human CRMs in our dataset. It has been previously reported that such enrichments tend to associate with CRMs carrying tissue-specific epigenetic signatures [32]. However, given the limited correlation between chromatin states and grammatical patterns we observed, we expected to find enrichments among both shared and tissue-specific grammatical patterns, though perhaps most strongly within the most deeply conserved patterns: the MCKG grammatical class. We tested this by intersecting human CRMs with variants from the NHGRI-EBI GWAS Catalog [33] and testing for significant enrichments of GWAS SNPs and associated GWAS ontology terms [34]. As expected, we observed a 1.2-fold enrichment of GWAS SNPs in all categories, (p < 4.4e-11), and a stronger, 1.5-fold enrichment for immune-specific GWAS SNPs (p < 5.9e-8) in CRMs relative to matched background sequences.

To evaluate the likelihood that CRM-associated SNPs serve causal roles, we compared the proportion of SNPs in CRMs and matched background sequences falling into each RegulomeDB class [35]. Overall, CRM-associated GWAS SNPs had systematically higher levels of functional evidence than did background-associated counterparts (Fig. 6B, Wilcoxon p-value < 3.0e-20). CRM-associated SNPs were 1.9 times more likely to fall into classes 1 and 2, which indicate high likelihood of affecting TF binding, compared to those within background regions (Fisher’s exact p-value 3.2e-22). By contrast, only 4% of CRM-associated SNPs, compared to 56% of background SNPs, fell into classes 5-7, for which there is minimal/no evidence for effects on TF binding. These data indicate a high likelihood for enrichment of causal SNPs among CRM-associated GWAS SNPs.

**Figure 6:**
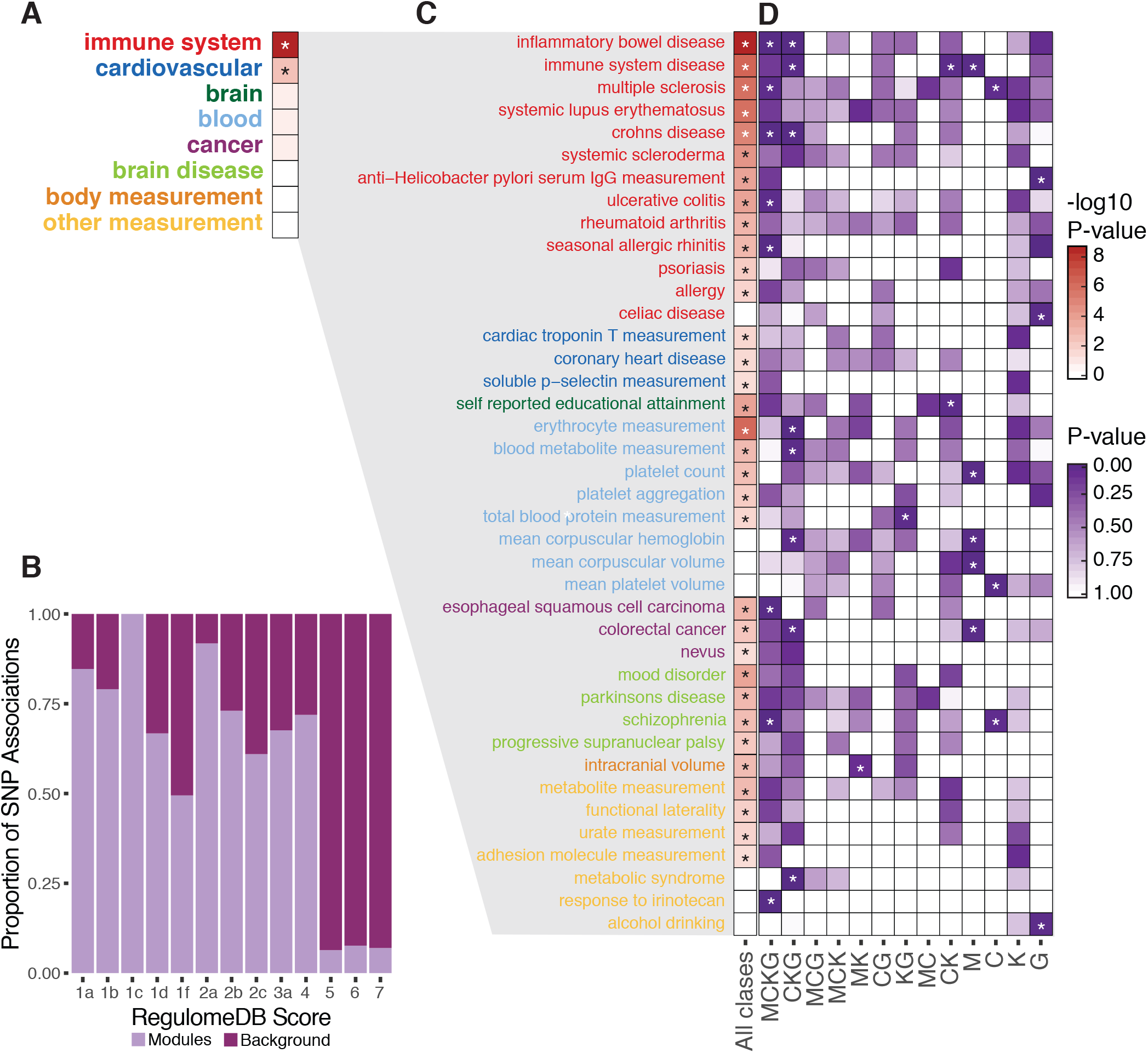
Grammatical patterns are enriched for relevant human GWAS causal SNPs. **A.** Fisher’s exact test p-values for enrichment of functional categories among GWAS SNPs in CRMs compared to matched background regions. P-values are shown for all categories in which at least one specific ontology term was enriched among CRMs in individual tests. Only two terms, “immune system” and “cardiovascular”, were significantly enriched. **B**. RegulomeDB scores were tabulated for GWAS SNPs in CRMs and matched background regions. The fractions of all observations from CRMs relative to background sequences within each RegulomeDB score category are shown, demonstrating marked enrichments of functional evidence among CRM-associated SNPs. **C**. Binomial enrichment p-values for GWAS ontology terms among CRM-associated GWAS SNPs. **D**. Binomial enrichment p-values for GWAS ontology terms among CRM-associated SNPs, separated by the grammatical class of the host CRM. Within all panels, stars indicate multiple-testing-corrected p-values <= 0.05.

To gain further insight into the pathways affected by these SNPs, we looked for enrichments of 27 categories of ontology terms, and for individual ontology terms (table s5). The only enriched categories were immune system (1.4-fold, p < 2.3e-90) and cardiovascular (1.2-fold, p < 2.8e-3) (Fig. 6A). In addition, we detected 33 individual ontology terms that were significantly enriched among CRM-associated SNPs across all grammatical classes (Fig. 6C). Of these, 12 were immune related, 10 of which ranked among the top fifteen terms, with a mean five-fold change over their expected binomial frequency (table S6). We also found seven terms that were significant within individual grammatical classes but not in the overall dataset (Fig. 6D, table S6). Notably, the majority of enriched ontology terms relate either directly to immunity, autoimmune disorders, or to cardiovascular disease, psychiatric disorders, and cancer, all of which have an underlying basis in, or known correlations with immune system dysfunction [36-42]. We also noted substantial overlaps between functional categories in our enriched in our dataset and those reported previously for the same cells [20]. It is interesting to note that our top term, “Inflamatory Bowel Disease”, was also significantly enriched in occupancy conserved TFBSs in GM12878 cells in Cheng et al. We observed strong enrichments for this term among MCKG and CKG grammatical patterns, although its enrichment in GM12878-specific patterns failed to reach significance.

Among individual grammatical classes, the MCKG and CKG classes were most likely to carry significant GWAS enrichments (Fig. 6D), indicating that GWAS SNPs disproportionately affect conserved regulatory logic. However, significant enrichments were also observed in GM12878-specific (G), Human-specific (KG), Myeloid-specific (MK), and two-cell-mixed (CK) grammatical classes. Surprisingly, we also see significant enrichments for 8 terms in the mouse-specific M and C classes. It is unclear how these regulatory patterns might influence the etiology of human disease. Nevertheless, the fact that the majority of GWAS SNPs affect conserved grammatical patterns suggests their importance in maintaining proper regulatory control of disease-related pathways. Accordingly, it is possible that enrichment for these patterns may indicate pathways for which regulatory control is conserved between human and mouse and, as such, may represent promising targets for translational research. Conversely, given the evidence we see for enrichment of species-specific grammatical patterns near genes with divergent expression, these patterns may highlight pathways for which regulation has significantly diverged in mouse, making them less promising as models for human disease.

### Human-Mouse SOM Data Browser

To facilitate data analysis and visualization, the SOM data are presented in a public browser with extensive search and visualization capabilities (https://boylelab.med.umich.edu/SOMbrowser/). Clickable density maps of various annotations are available for visual comparisons between patterns, with CRM-level annotations for the selected pattern given in a tabbed browsing pane, including relevant links to external resources to facilitate in-depth analysis. Two additional tools, the “compare maps” section, and a flexible search tool, are available to facilitate further comparisons and customized views of the data. The compare maps tool allows side by side comparison of multiple, user-selected maps, while the search utility allows users to construct arbitrarily complex queries directly against the browser database. Search results are presented graphically, in a map that shows the density of results in each grammatical pattern, alongside detailed tabular results. Both sections offer direct links to the main browser for further analysis. Integrated help links are provided throughout the browser, as well as a detailed help section with instructions on how to use each feature.

## DISCUSSION

The results we present build upon previous significant studies in this area [7,14,17,21-23,25] by expanding our understanding of the relationship between positional conservation and underlying gene regulatory logic. We have shown widespread differences in the specific regulatory logic occupying positionally-conserved loci across species and cell types, and corresponding differences in underlying chromatin states, with positional conservation being a poor predictor for shared chromatin states between species and cell types.

By contrast, grammatical patterns have stable functional signatures that span both species and tissue, and these signatures remain relatively constant regardless of the evolutionary history of CRMs. This property gives grammatical patterns a distinct advantage for predicting the function of regulatory sequences compared to methods relying on positional conservation of TF occupancy and/or other functional annotations, given that we know a large fraction of regulatory sequence does not map between even closely-related species. This is of particular importance given that transposable element insertions have been shown to propagate TFBSs within multiple mammalian genomes, and these insertions have been shown to directly modify certain regulatory functions in the genome, including formation and/or maintenance of chromatin loops [2]. Our results show that species-specific CRMs share the same functional signatures as positionally-conserved CRMs within their grammatical pattern. Surprisingly, we show that these species-specific CRMs also overwhelmingly carry conserved regulatory logic.

We also show a clear correlation between the inherent tissue/species-specificity of grammatical patterns, expressed by their grammatical class, with matching tissue/species-specific gene expression patterns. This not only supports our theory that grammatical patterns are predictive of tissue/species-specific regulatory activities, but also suggests that grammatical classes capture the functional conservation of regulatory logic across tissues and species. As such, patterns specific to a given tissue may offer unique opportunities to understand the regulatory basis of relevant diseases, especially when these patterns are shared across species. By contrast, grammatical patterns specific to a single cell type may highlight disease-related regulatory mechanisms that have diverged between species. Similarly, species-specific grammatical patterns may represent emergent portions of the regulatory language with potentially causal roles in evolutionary divergence and speciation.

We show that grammatical patterns are significantly enriched for causal GWAS SNPs relevant to immune phenotypes, and that these enrichments fall both in shared and species-specific regulatory logic. It is interesting to note that we found relevant enrichments not only among CRMs with human-specific grammatical patterns, but also in human CRMs labelled with mouse-specific grammatical patterns. While it is not immediately clear what might explain this phenomenon, one appealing hypothesis is that these CRMs represent inappropriate, out-of-context use of these regulatory instructions which may, in itself, lead to a diseased state. In this framework, the underlying GWAS SNPs may directly cause the grammatical pattern to shift from a human pattern to a mouse pattern by creating or ablating a TFBS, thus leading to target gene dysregulation. Indeed, the fact that these experiments were run using immortalized cell lines may increase the likelihood of observing such cases.

While previous studies have used shared occupancy across species as a proxy for shared regulatory logic, we find that the functional annotations used in this study correlate better with grammatical patterns than conserved occupancy, with the notable exception of PhastCons conservation. We find that the segmentation of TF co-binding patterns with SOMs allows a much clearer picture of underlying regulatory potential and conservation than does looking at the regulatory landscape through a purely phylogenetic lens. SOMs have been used previously to study gene regulation, in a study of TFBS co-binding profiles across multiple human cell types [7], across distant species [14], and, most recently, to investigate combinatorial regulatory logic in the mouse liver [43]. However, as far as we know, this is the first application of these methods to groups of cells spanning more than one mammalian species.

Although the analysis presented by Dubois-Chevalier et al. was referenced specifically to the liver-specific TF NR1H4, and contains few overlapping TFs with our dataset, we note important parallels between their results and ours. Primary among these are the ability of the SOM to separate grammatical patterns (termed trans-regulatory modules, or TRMs, in their paper) into clusters with chromatin signatures characteristic of enhancers or promoters, and the association of distinct sets of TRMs with expression (and dysregulation) of liver-specific genes. We observed both of these characteristics in our dataset as well, in the predictive value of grammatical patterns for underlying chromatin states, and in enrichments of tissue/species-specific grammatical patterns near genes with corresponding expression.

Another group has applied SOMs to classify regulatory sequences into cell-specific and shared functional groups across a number of human cell types based on histone modifications, including those used in this study [32]. They went on to map binding data and motifs for several general and tissue-specific TFs to their dataset, and found that tissue-specific sets of histone modifications were significantly correlated with presence of corresponding tissue-specific TF motifs and ChIP-seq peaks [32]. While it is tempting to relate these data to our, seemingly contradictory, finding that matched chromatin states have little predictive value for underlying TF binding, such a comparison may be misleading for two reasons. First, Mortazavi et al. found that tissue-specific epigenetic signatures predicted only presence of tissue-specific TFs and motifs. Unfortunately, our dataset does not contain appropriate tissue-specific TFs on which we could base a comparison. Second, our results are based on genome annotation using a seven-state chromHMM model. As a result, the genome segmentation used in our analysis is much coarser than the 1350-unit SOM on which their results are based. It is possible that, by dissecting the epigenetic landscape in such detail, one can find subsets of co-occurring modification that segregate distinctly with specific sets of co-bound TFs. However, their failure to find specific enrichments of any TFs within SOM units shared between all cells suggests that even highly detailed epigenetic annotations may not accurately predict underlying regulatory logic in many cases.

We must note that the focus of this study on only two sets of matched cell types with immune functions, owing solely to the availability of sufficient ChIP-seq data in the ENCODE repository at the time of the analysis, may restrict how broadly these specific results can be interpreted. Similarly, the use of immortalized cell lines may have introduced confounding factors, as it is currently not known how cancer affects the regulatory lexicon as cells transition from a normal state to a diseased state. However, it is encouraging that we found such a strong overlap between regulatory patterns present across all four cells, which suggests that most of the regulatory landscape has remained intact despite the many other genomic and physiological changes these cell lines harbor. However, another interpretation would be that all cancer cells carry similar changes in regulatory grammar, and we cannot rule this out using the current dataset.

## CONCLUSIONS

On the whole, these data demonstrate that grammatical patterns can augment and extend our ability to classify and predict regulatory functions beyond use of positional conservation. The SOM allows us to segment the regulatory landscape into grammatical patterns that may be shared among all, or specific to any subset of, species and cells used in map construction. We have shown that these grammatical patterns are predictive of underlying functional annotations, correlate with species-specific gene expression patterns, and are enriched for GWAS SNPs for relevant phenotypes. The marked enrichment of GWAS SNPs within highly-conserved grammatical patterns further suggests that they may be a useful tool to identify disease mechanisms and pathways best suited for translational research. We believe that, as data become available in a broader array of primary and immortalized cell types, we can gain great insight into both shared and divergent portions of regulatory grammar between human and mouse. This may shed further light on why mouse models often fail to translate to the human system, and highlight pathways that are most likely conserved in their regulation between both species.

## Methods

### Preparation of the Dataset

Data for the analyses performed in this manuscript are available from the ENCODE project (https://www.encodeproject.org/). ChIP-seq experiments follow ENCODE standards for preprocessing, quality control and uniform peak calling. All data were downloaded from the ENCODE DCC portal. Human data map to hg19 and mouse data to mm9. Data used and accessions are available in table S1.

We collected 27 ChIP-seq datasets for which there were data in both human K562 and GM12878 as well as data for their 1-to-1 orthologs in mouse MEL and CH12 cell types. All regions in each species were merged using BedTools [44] mergeBed to create a base set of cis-regulatory modules (CRMs) in each species. These regions were then labeled with TF binding data specific to each cell type. Only regions with at least two TFs bound were considered for further analysis.

The SOM was constructed using methods defined by us previously. Briefly, we train a 47x34 toroidal SOM with hexagonal neurons with an update radius of ⅓ map size and alpha of 0.05 for 100 epochs. We repeat this for 1000 trials to minimize the quantization error. We perform this analysis with a modified form of the kohonen R package available through our github site here: https://github.com/Boyle-Lab/kohonen2

In contrast to our previous methods, we also performed 10,000 random permutations of the SOM using the same set of parameters. We permuted labels of TFs on each region but kept cell type and organism constant for those labels. This allowed for us to correct for patterns that occur at low frequency by chance and for factors with many apparent associations due to their high frequency in the datasets. Using this empirical distribution we considered a neuron significant if it had an empirical p-value <= 0.0001 (i.e. has not appeared in any random permutations). In order to avoid thresholding issues, we then only considered differential regions to be those with an empirical p-value > 0.95 in non-enriched cell types.

Each hexagonal neuron was assigned a corresponding grammatical pattern by enumerating occurrences of each of the 27 TFs in all CRMs assigned to a given neuron. If a TF occurred in at least 90% of CRMs assigned to a neuron, it was considered to be an integral part of the grammatical pattern. Similarly, grammatical classes were assigned using the empirical p-values calculated for each cell, requiring an empirical p-value <= 0.0001 to include a cell type in a pattern’s grammatical class and a p-value > 0.95 to exclude a cell from a pattern’s grammatical class. Putative grammatical patterns where not all cell types could be assigned as conserved or differential according these upper and lower p-value thresholds were considered ambiguous in their grammatical conservation and were excluded from the dataset.

### Preparation of Background Sequences

To serve as a baseline for our statistical tests, we randomly selected sets of background regions, matched in length, GC content, and distance to the nearest TSS, from both the human and mouse genomes. These were analyzed following the same procedures as CRMs whenever possible. Briefly, CRMs were selected at random from the human or mouse dataset, without replacement, and the length, GC content and distance from the midpoint of the element to the nearest TSS were calculated and assigned to one of five bins: 0-500bp, 501-2,000bp, 2,001-5,000bp, 5,001-10,000bp, and >10,000bp. A maximum of 1000 sequences of matching length were randomly drawn from the genome, excluding the CRMs themselves, until a sequence with GC content +/- 10% and TSS distance within the same distance bin was found. This sequence was labeled with all metadata from its source module, stored within the background dataset and masked from the genomic fasta to prohibit redrawing the same sequence. This process was repeated until either a) all CRMs were exhausted, b) all background windows were exhausted, or c) 100,000 background sequences were drawn. For quality control, density plots of the distributions of length, GC content and TSS distance were superimposed and visually assessed, and means were compared using the t.test function in R. No significant differences in these parameters were found between the source and background distributions.

### Database and Browser Preparation

All data pertaining to the patterns and CRMs in the dataset, including background sequences and annotations resulting from all analyses performed, were loaded into a mysql database to enable further data dissection and intersection and serve as the data repository for the SOM data browser. Data were retrieved through a combination of manual sql queries and automated access through the mysql PERL API. The SOM Browser web application was built around the open source Catalyst framework (http://www.catalystframework.org/) and interfaces directly with the database for visual display and analysis through various precomputed data views and an extensible search utility.

### Analysis of orthologous and non-orthologous sequences

In order to investigate the interspecies relationships between the 84,713 human and 125,055 mouse CRMs, we used bnMapper [24] to cross-map CRMs and background sequences between genomes using settings described in [20]. Human-to-mouse and mouse-to-human liftover chains were obtained from the UCSC download portal (http://hgdownload.soe.ucsc.edu/downloads.html) [45]. One-to-one orthology is not enforced in these sets of chains, which we will call “one-to-many” chains. We also produced reciprocal-best chains, representing the single most likely set of one-to-one orthologs between both species, as described at (http://genomewiki.ucsc.edu/index.php/HowTo:_Syntenic_Net_or_Reciprocal_Best). We used the reciprocal-best chains to map putative 1:1 orthologs between species, which are used as our benchmark “orthologs” dataset. Non-orthologs were identified by excluding all CRMs mappable using the one-to-many chains from the original dataset. CRMs that mapped using the one-to-many chains but not the reciprocal-best chains were considered likely one-to-many orthologs, and these were excluded from further analysis, as were a small number of CRMs that mapped using the reciprocal-best chains but not the one-to-many chains. CRMs were assigned to a positional class based on the cell(s) in which we observe ChIP-seq occupancy within the same genomic interval and/or an overlapping orthologous region. CRMs and background sequences were assigned numeric codes: 0 (non-ortholog), 1 (1:1 ortholog), or 2 (one-to-many ortholog), which were used to classify and filter the datasets in later analyses.

### Analysis of module gain and loss

Starting with all CRMs labeled as strict non-orthologs (orthology code 0) in our initial mapping step, we employed a phylogenetic maximum parsimony method to infer whether each non-orthologous element had been gained on the branch leading to its host species or lost on the branch leading to the species in which it is absent. We first retrieved the 46-species placental mammal phylogenetic model from UCSC (http://hgdownload.cse.ucsc.edu/goldenpath/hg19/phyloP46way/placentalMammals.mod)[45]. The tree_doctor utility from the Phast package [46] was used to prune all branches but human, mouse and three outgroups: dog, horse and elephant. Liftover chains for human and mouse to each of the outgroup species were obtained from the UCSC download portal [45] and bnMapper [24] was used to cross-map human and mouse sequences to each outgroup, using the same methods and settings described in the “Analysis of orthologous and non-orthologous sequences” section. Given these mappings, the ancestral state at the most recent common ancestor (MRCA) was inferred by maximum parsimony using a custom perl script. In cases where the ancestral state could not be unambiguously resolved using the whole phylogeny (roughly 12% of all unmapped CRMs), we dropped the terminal outgroup, elephant, from the phylogeny. This allowed us to unambiguously assign 100% of unmapped CRMs and background sequences as either gains or losses on a specific branch. Numeric orthology codes were reassigned for these elements in the database: 3 = mouse gain, 4 = human gain, 5 = mouse loss, 6 = human loss. These codes were used to extract CRMs in each orthology class for later steps in the analysis.

### Analysis of Conservation

PhastCons conserved elements and base-wise scores for the human (46-way placental mammals) and mouse (30-way placental mammals) genomes were retrieved from the UCSC download portal in bed format [45]. Intersections and coverages of CRMs and background sequences by phastCons elements were extracted with bedtools intersect and bedtools coverage [44] and the coverage, average, maximum and minimum scores for each module were extracted using a custom perl script. Bulk enrichments over background were tested in R using the fisher.test function with default options and p-values were corrected for multiple testing by the holm method. Data were loaded into the database and SQL queries were used to extract intersections of CRMs and matched background regions with PhastCons elements for individual grammatical patterns and six aggregate grammatical classes: Human Specific (K, G, KG), Mouse Specific (M, C, MC), Lineage Specific (MK, CG), Two-Cell Mixed (MG, CK), Three Cell (MCG, MCK, MKG, CKG), and MCKG, within orthologous, non-orthologous, gain and loss fractions for both human and mouse. These were stored in text files and loaded into R in order to test each subset for enrichment over background. Fisher’s exact tests were performed with alternative “greater” and multiple testing correction was performed with the holm method. Percent intersections of CRMs with phastCons elements for all subsets were plotted using the ggplot2 R package (Fig. 3C).

Heat plots of PhastCons conservation scores presented in fig S4 were produced by extracting PhastCons score vectors from the hg19 phastCons46way, and mm9 phastCons30way bigWig files, retrieved from the UCSC download portal [47], with a custom R script based on the RTrackLayer package. We first randomly selected 1000 ChIP-seq peaks from ortholgous, loss, and gain CRMs, from both human and mouse. PhastCons score vectors were retrieved for 400 bp windows centered over each ChIP-seq peak summit, using our custom function, and stored as matrix rows. Matrices were then sorted by descending row sum and plotted using a custom plotting function in R.

### Analysis of Histone Modifications

ChromHMM chromatin state predictions were obtained from the ENCODE DCC portal for all cell types (table S1). All chromatin state annotations intersecting CRMs and background sequences and their orthologous locations in human/mouse were extracted from the bigBed files with a perl script and loaded into a database table. Subsets of the data for each aggregate grammatical class were then extracted from the database from the command line and stored in text files for later loading into R, or directly in R using the RMySQL package, for further analysis. We focused on four chromatin states annotated in the chromHMM data: promoter (H3K4me3), strong enhancer (H3K4me1+H3K4me3), weak enhancer (H3K4me1), and polycomb repressed (H3K27me3).

In order to test for enrichments of ACS in CRMs relative to matched background sequences, we used SQL queries to extract intersections with ACS for the six aggregate grammatical classes described in the “Analysis of Conservation” section, for both CRMs and background sequences. The resulting matrix of counts was read into R and Fisher’s Exact tests for enrichment of ACS in CRMs over background were performed (table s4). Percent intersections with ACS were plotted with the ggplot2 R package (Fig. 3B).

We quantified the predictive value of grammatical patterns regarding underlying chromatin states by comparing the chromatin state annotations for all pairs of loci within each grammatical pattern, for promoter, strong enhancer and weak enhancer states (Fig 4A, fig S5B). Similarly, we quantified the extent to which promoter, strong enhancer, and weak enhancer chromatin states were predictive of the underlying regulatory grammar at the same/orthologous locus across cells and species by counting the fraction of sequence pairs with matching chromatin states at each locus in the dataset that also match in grammatical pattern, after excluding two major cohesin-related grammatical patterns (Fig 4B, fig S5A). Euler diagrams in Fig 4 and fig S5 were generated using the draw.pairwise.venn function from the VennDiagram R package.

Evaluation procedures for cell-specificity of ACS within grammatical patterns are shown in fig S6A, for the summary hat map presented in Fig 4C, and s6C, for the full heat map presented in fig S7A. Counting procedures for the full matrix (fig S7A) are described in fig S6B. Iterating over each combination of chromatin state and cell, we selected all grammatical patterns for which at least 50% of the CRMs from the reference cell were annotated with a given chromatin state and counted the occurrences of the same mark in each of the other three cell types to produce the matrix of counts overlayed on the heat map in fig s7A. These were visualized as a heat maps using the geom_tile function from the ggplot2 R package (fig S7A), with the density scale fixed to a range of (0, 1) to enable direct comparison of maps. The summarized heat map in Fig 4C were prepared by defining “within class” cells as those that are included in the given grammatical class (e.g., MEL and CH12 for the “MC”), and “outside class” cells as those not included in the given grammatical class (e.g., K562 and GM12878 for “MC”). Counting procedures are described in fig S6A. For all combinations of grammatical class and chromatin state, the number of grammatical patterns in which >= 50% of CRMs was recorded in the “total” column of the 3x3 count matrix. The “in class” column was counted for each row as the number of patterns comprising the “total” column in which >= 50% of CRMs from cells within the assigned grammatical class intersect the matching chromatin state. The “outside class” column was calculated as the number of patterns comprising the “total” column that intersect the given chromatin state in >= 50% of CRMs from cells outside the corresponding grammatical class.

Evaluation procedures for cell-specificity of ACS within positional classes are illustrated in fig S6B, for the summary matrix (Fig 4D), and the full matrix (Fig S7B) in fig S6D. For the summary matrix (Fig 4D), we defined three classes of loci: matches, in which the same histone marks are found in all occupied cell types and no unoccupied cell types; mismatch 1, in which a locus carries the same histone marks in all occupied cell types and one or more unoccupied cell types; and mismatch 2, in which not all occupied cell types carry the given histone mark. The resulting 3x4 matrix of positional locus counts were expressed as heat maps with geom_tile and a fixed shading scale as before. For the full matrix (fig S7B), iterating over all combinations of chromatin state, cell, and positional class, we counted the number of CRMs from the reference cell marked with the given chromatin state, and, for each of those loci, the number of times the same state was observed in each of the three other cells. The resulting count matrix was visualized as described for grammatical classes.

Graphical plots of chromatin state intersections for grammatical pattern 821 (fig S5C) were prepared by extracting the chromatin state annotations spanning the length of all CRMs within pattern 821 from the database through the RMySQL package. The annotations within each CRM were scaled and sorted such that all states present in a given CRM are shown in ascending numerical order along the length of the plot row, as different colors, with the length of each color corresponding to the fraction of the CRM intersecting the given state. Plot rows were then sorted by descending row sums and visualized using a custom plotting function. Plots of chromatin states for CG positional class loci in Fig 4D were produced in similar fashion, except that state intersections for all occupied and unoccupied cells in each of the 10 loci shown were extracted from the database, followed by within-row sorting and plotting of locus-wise groups using a custom plot function.

### DNase Hypersensitivity Analysis

DNaseI hypersensitivity hotspots were retrieved from the ENCODE DCC portal in bed format (table S1). All datasets within each cell type were combined, sorted and merged with bedtools merge prior to calculation of intersections and coverages with bedtools intersect and bedtools coverage [44]. Cell-wise coverages for all CRMs, background sequences and their orthologous locations, if present, were compiled into a table with a custom perl script, and loaded into the database. SQL queries were used to fractionate the data into human and mouse subsets of orthologous, nonorthologous, gain and loss classes for CRMs and background sequences in each of six grammatical categories: human-specific, mouse-specific, lineage-specific (myeloid/lymphoid), two-cell-mixed, three-cell and ubiquitous (MCKG). These counts were assembled into tables and read into R for Fisher’s Exact tests as described in the “Analysis of Conservation” section. Plots of the percentage of CRMs overlapping DNase hypersensitive elements in each subset were constructed in R using the ggplot2 R package (Fig. 3A).

### Analysis of Target Gene Expression

Single-ended RNA-seq sequencing data for human K562 and GM12878, and mouse MEL and CH12 cells were obtained from the ENCODE download portal in fastq format (table S1). In order to mitigate the impact of sequencing batch effects, whenever possible, we restricted our choice of datasets to those produced within the same laboratory using the same protocols and sequencing equipment. Kallisto version 0.42.4 [48] was used to quantify expression levels directly from the fastq sequencing data without an intermediate mapping step. Initally, genomic sequences for all annotated transcripts in the UCSC “knownGene” tables for hg19 and mm9 [45], were extracted from the human and mouse genomes as fasta sequences, using gffread 0.9.5 (https://github.com/gpertea/gffread). Transcript indexes were prepared from these files using the “kallisto index” command. We then used “kallisto quant”, with options -b 100 -single -l 200 -s 80 -t 8, to calculate raw expression values and bootstrap data for each replicate individually. Replicates were combined and expression values normalized within each species using the sleuth R package (http://pachterlab.github.io/sleuth/). R analysis scripts are available through our github repository (https://github.com/Boyle-Lab/mouse-human-SOM). UCSC transcript identifiers were converted to gene names with a custom perl script, based on a translation table obtained from the UCSC Table Browser [49] and gene-wise TPM values were calculated by totaling the expression values for all isoforms corresponding to a given gene. These were assembled into a 6xN table of gene symbols for both species for all genes with 1:1 orthologs, or a gene symbol and null placeholder for species-specific genes, and associated expression values in each cell. Gene ortholog relationships were based on the modencode common orthologs list [50], as obtained from (http://compbio.mit.edu/modencode/orthologs/modencode.common.orth.txt.gz). This table was read into an R matrix and quantile normalized using the normalize.quantiles function from the preprocessCore package.

As an alternative method to assess gene expression levels, we obtained the bam alignment files corresponding to the same fastq datasets described previously and performed an explicit correction process for batch effects, as described in [51]. Initially, UCSC knownGenes transcripts were collapsed into a gene-centric gff file using the mergeOverlappingExons.py script published in [51]. Gene-wise GC contents were computed using bedtools nuc [44] and raw fragment counts were extracted from the bam files with featureCounts [52] based on this gene-wise gff. Counts were totaled over both replicates for each cell type and assembled into a 4xN matrix, with N being the number of genes annotated in the dataset. Counts were read into R and normalized following the exact process described in [51]. To obtain a more complete set of expression values, we also performed normalization with one modification: we did not filter out the bottom 30% of genes by expression value. We compared the resulting datasets by overlaying density plots of the expression values for genes in both datasets and by performing a paired t-test. Density plots were visually indistinguishable and the mean difference observed in the t-test was very small (0.016), although with a significant p-value (5.987e-15), owing to the large size of the dataset. We used this complete expression dataset for all analyses. Finally, we converted the normalized counts to TPM to facilitate direct comparison to the Kallisto expression values, using the weighted average of the all the transcript-wise effective lengths for each gene, as reported by Kallisto, as the effective length of the gene.

Differential gene expression analysis was performed with DESeq [31]. Raw counts for all RNA-seq replicates were first normalized for batch effects following the procedures described above. DESeq size factors were fixed at 1 to disable internal normalization and differential expression predictions were performed using the nbinomGLMTest procedure using a full model including both species and cell effects and a reduced model including only species. Predictions with multiple-testing-corrected p-values <= 0.05 were retained as differentially-expressed. From these, we selected gene sets with expression specific to a given species or cell type based on the predicted partial slope coefficients in the full model. For example, myeloid specific genes were selected by setting a lower-bound of 4 on the "cellTypeMyeloid coefficient and upper, indicating a positive correlation with expression in myeloid cells, and lower bounds of 0.05 and -0.05 on the "speciesMm9 coefficient, indicating little or no correlation with species. Similarly, a set of stably expressed genes was selected from among all predictions by setting the upper and lower bounds for both coefficients at 0.065 and -0.065, yielding a set of 724 stably-expressed genes.

Nearest-gene assignments for CRMs were made based on the nearest TSS, obtained from another analysis using Chip-Enrich [53]. Counts of CRMs in each grammatical class nearest to genes in each set were retrieved from using SQL queries, directly into R tables using the RMySQL package. Within each aggregate grammatical classes, we calculated the fractions of CRMs nearest to genes with stable expression (71S) and CRMs nearest to in-class genes (71D) and calculated a tissue-specificity coefficient, *c*, as *c* = log2(πD / πS), where *c* > 0 indicates tissue-specific enrichment and *c* < 0 indicates tissue-specific depletion. We also calculated a corresponding coefficient, c′, as c′ = log2(πD′ / πS), where (πD′) represents the fraction of CRMs nearest to cross-class genes. For comparing the cell-specificity coefficient frequency of CRMs within each aggregate grammatical class inside their corresponding gene set and the stably-expressed set. We visualized these log-ratios as pairs of bars in Fig 5, produced using the ggplot2 geom_bar function, interpreting positive values as evidence for enrichment, and negative values as evidence for depletion. Pairwise significance tests were performed using the fisher.test function in R with alternative “greater”, for tests of enrichment among CRMs nearest genes with matching specificity, or “less”, for tests of depletion of CRMs nearest genes with mismatched specificity.

### Analysis of Genome-Wide Association Study SNPs in Human

Initially, a list of human GWAS lead SNPs, in hg38 coordinates, and their associated GWAS Ontology terms [33] was obtained from the NHGRI-EBI GWAS Catalog web site (https://www.ebi.ac.uk/gwas/). Coordinates for GWAS Catalog SNPs were converted to hg19 by matching to dbSNP build 142 based on RSID. 61 records that initially failed to map based on RSID were found to have been merged into other dbSNP records and were successfully mapped using the active RSID. The 46 remaining unmapped SNPs were mapped to hg19 using liftOver [54]. During this process, four GWAS Catalog records were found to contain anomalies in the RSID field: one with a space character inserted (rs1083 2417), which was resolved by removing the space, and three with nonnumeric characters appended (rs4460079b, rs7658266b, rs6917824L), none of which could be found in the cited references under the given RSID, minus the trailing letter, or any alternate RSIDs in dbSNP. These three records were excluded from the analysis. We next found all SNPs in linkage disequilibrium with the hg19-referenced GWAS SNPs at R^2^ >= 0.8, based on hapMap linked SNP data, using a custom perl script. These were merged into the dataset, which was subsequently intersected with the CRMs and background sequences with bedtools intersect [44]. CRMs containing GWAS SNPs and their associated GWAS Ontology terms were counted using a combination of command-line utilities and custom perl scripts. Duplicate GWAS Ontology terms within a single module were counted only once. GWAS Ontology terms were categorized into 29 groups (table S5) based on the EBI Experimental Factor Ontology and primary literature sources, describing known associations with specific tissues, disease states, biological functions and processes similar to [21]. Single-tailed Fisher’s Exact tests were used to assess enrichment of GWAS SNPs as a whole and for each of the 29 functional categories in CRMs relative to background sequences. Enrichments of individual GWAS Ontology terms in CRMs were tested using binomial tests with the binomial probability set to the relative frequency of each term within the GWAS Catalog. All tests were performed in R and corrected for multiple testing using the Benjamini-Hochberg method.

RegulomeDB scores for all SNPs within CRM and background datasets were extracted from the RegulomeDB web site using an automated scraper, written in Python, and read into R. CRM and background score distributions were compared using the wilcox.test function with alternative = “lower”. Frequency tables of RegulomeDB scores were produced and normalized to the total number of observations within the CRM and background datasets before visualization with ggplot.

## DECLARATIONS

### Ethics approval and consent to participate

Not applicable.

### Consent for publication

Not applicable.

### Availability of data and material

All analyses available through https://boylelab.med.umich.edu/SOMbrowser/ and https://github.com/Boyle-Lab/mouse-human-SOM. All data are available through the ENCODE project at <u>encodeproject.org</u>.

## Acknowledgements

We thank the ENCODE project for making well-documented data freely available for use in this project.

